# Broad-Spectrum Antifungal Activities and Mechanism of Drimane Sesquiterpenoids

**DOI:** 10.1101/816082

**Authors:** Edruce Edouarzin, Connor Horn, Anuja Paduyal, Cunli Zhang, Jianyu Lu, Zongbo Tong, Guri Giaever, Corey Nislow, Raja Veerapandian, Duy H. Hua, Govindsamy Vediyappan

## Abstract

Eight drimane sesquiterpenoids including (-)-drimenol and (+)-albicanol were synthesized from (+)-sclareolide and evaluated for their antifungal activities. Three compounds, (-)-drimenol, (+)-albicanol, and (1*R*,2*R*,4a*S*,8a*S*)-2-hydroxy-2,5,5,8a-tetramethyl-decahydronaphthalene-1-carbaldehyde (**4**) showed strong activity against *C. albicans*. (-)-Drimenol, the strongest inhibitor of the three, (at concentrations of 8 – 64 μg/ml, causing 100% death of fungi), acts not only against *C. albicans* as a fungicidal manner, but also inhibits other fungi such as *Aspergillus*, *Cryptococcus*, *Pneumocystis*, *Blastomyces*, *Fusarium*, *Rhizopus*, *Saksenaea* and FLU resistant strains of *C. albicans*, *C. glabrata, C. krusei*, *C. parapsilosis* and *C. auris.* These observations suggest drimenol is a broad-spectrum antifungal agent. At high concentration (100 μg/ml), drimenol caused a rupture of the fungal cell wall/membrane. In a nematode model of *C. albicans* infection, drimenol rescued the worms from *C. albicans*-mediated death, indicating drimenol is tolerable and bioactive in a metazoan. Genome-wide fitness profiling assays of both *S. cerevisiae* (nonessential homozygous and essential heterozygous) and *C. albicans* (Tn-insertion mutants) collections revealed putative genes and pathways affected by drimenol. Using a *C. albicans* mutants spot assay, the Crk1 kinase associated gene products, Ret2, Cdc37, and novel putative targets orf19.759, orf19.1672, and orf19.4382 were revealed to be the potential targets of drimenol. Further, computational modeling results suggest possible modification of the structure of drimenol including the A ring for improving antifungal activity.

## INTRODUCTION

Life-threatening fungal infections are an important cause of morbidity and mortality, particularly for patients with immune deficiency and those who are undergoing chemotherapeutic treatments. Some of the leading invasive fungal pathogens include *Candida* sp., *Aspergillus* sp. and *Cryptococcus* sp. Currently, the antifungal therapeutic options are limited, especially when compared to available antibacterial agents (1–4). Among the five classes of antifungals, azoles, echiocandins, polyenes, allylamines, and pyrimidine derivatives, only three are used clinically; azoles, echiocandins, and polyenes. Azole drugs, such as fluconazole (FLU), inhibit ergosterol synthesis through inhibition of lanosterol 14-α-demethylase, impairing formation of the fungal cell wall. Echocandins, such as caspofungin (CAS), block 1,3-β-glucan synthase and lead to depletion of glucan in fungal cell wall. Polyenes, including amphotericin B (AMB), bind to ergosterol in fungal cell membrane and change the cell membrane transition temperature, resulting in leakage of ions and small organic molecules, and eventual cell death. Allylamines, such as amorolfin, affect ergosterol synthesis by inhibition of squalene epoxidase. Pyrimidines, such as flucytosine (or 5-fluorocytosine), block nucleic acid synthesis, leading to the inhibition of protein synthesis (5, 6). No new antifungal agents have been approved by Food and Drug Administration since 2006 (2) and the latest antimycotic agents, echinocandins, were developed over 30 years ago (7, 8). Thus, new broad-spectrum antifungal agents are sorely needed to overcome the increasing emergence of antifungal drug resistance.

*C. albicans* is the most frequently found fungal pathogen in humans and costs the US health care system around $3 billion annually in treatment and lost productivity. *C. albicans*, a polymorphic fungus, exists as yeast, pseudohyphal and hyphal forms, with each contributing to its virulence. While the yeast form is essential for dissemination, the hyphal form is critical for invasion of cells, immune evasion, and biofilm formation. Furthermore, the ability to switch between forms is also essential for pathogenicity. Traditional antimycotics have many drawbacks, including toxicity to human cells, a limited range of cellular targets, the development of antifungal resistance (9–12), and the failure to successfully control pathogenesis. To develop new antifungal agents based on drimane sesquiterpenes, we have investigated synthetic drimane terpenes, (-)- drimenol (**1**) and (+)-albicanol (**2**), along with six analogs, **3** – **8** (Fig. 1), for their antifungal activities and identified (-)-**1** as a potent broad-spectrum fungicidal agent. Moreover, we determined their mechanism of action through forward genetic screening of mutant libraries of *C. albicans* and baker’s yeast and found that (-)-**1** affects the fungal activities of protein secretion, vacuolar biogenesis, chromation remodeling and cyclin dependent protein kinases (CDK) activity.

**FIG. 1.**
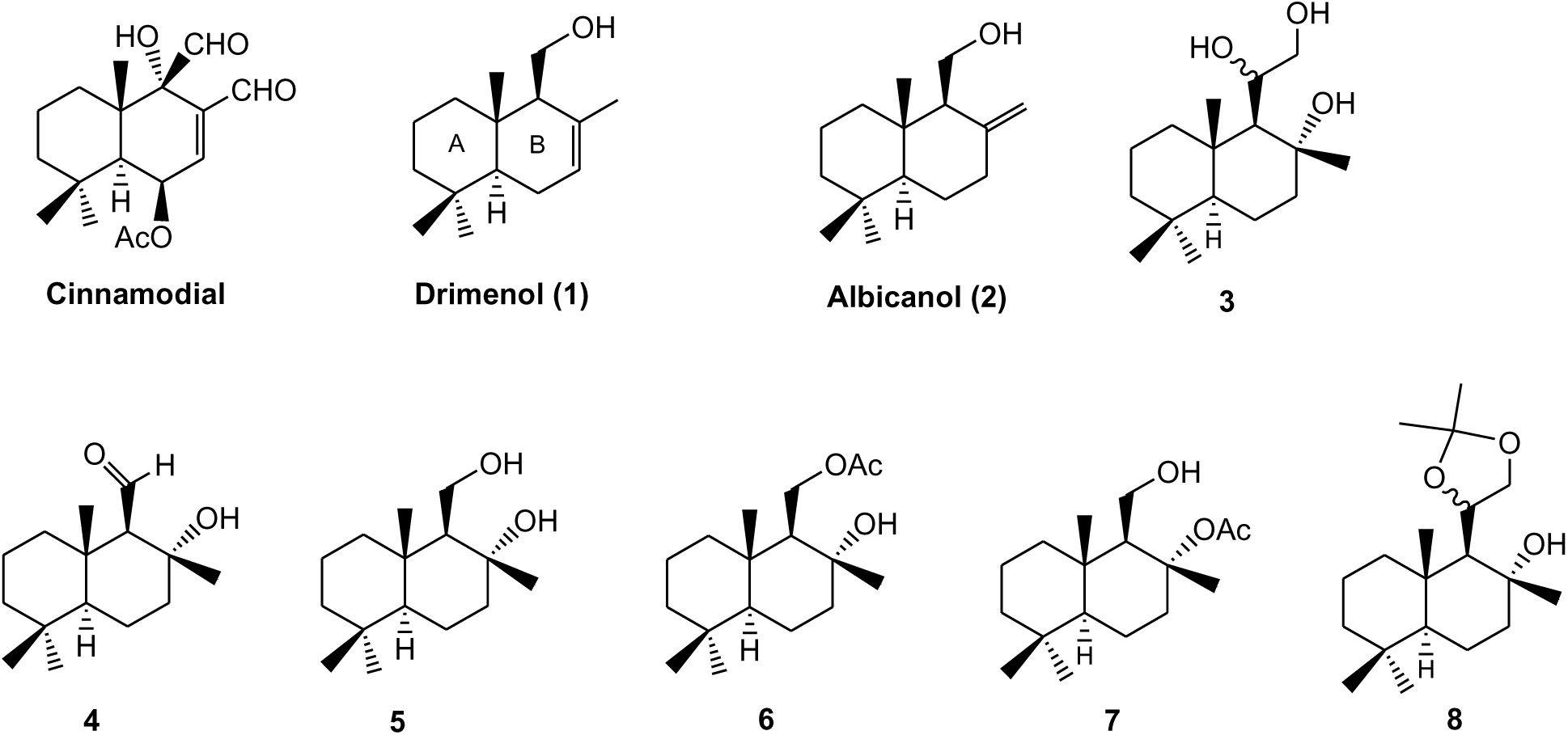
Drimane sesquiterpenoids bioevaluated for their antifungal activities.

## RESULTS

Several drimane sesquiterpenoids were synthesized in our laboratory during the total synthesis of (+)-chloropuupehenone, a natural product from marine sponges (13). Based on the antimycobacterial activity of sesquiterpene natural product, cinnamodial, isolated from the liverwort plant, *Warburgia salutaris* (14), we anticipated that drimane sesquiterpenes and closely related compounds (15) could be effective antimycotics for *C. albicans*. Five representative drimane terpenes, **1** – **5**, along with their derivatives, **6** – **8** (Fig. 1), were screened for their ability to inhibit *C. albicans* growth. It was assumed that additional hydroxyl group(s) or oxygen atoms in the molecule enhances water solubility and may improve bioactivity (16). Molecules **3** – **7** possess extra hydroxyl, aldehyde, or acetoxy function and molecule **8** contains an acetonide moiety in the drimane structure.

Compounds **1** – **5** were prepared by following previously reported methods (13). Molecule **6** was prepared from a mono-acetylation of **5** with acetic anhydride and pyridine in dichloromethane (Scheme 1). Molecule **7** was made by a sequence of three reactions: (i) silylation of the primary alcohol with *t*-butyldimethylsilyl chloride and imidazole in dichloromethane; (ii) acetylation of the tertiary alcohol with acetyl chloride and pyridine; and (iii) removal of the silyl ether group with tetra-*n*-butylammonium fluoride in THF. Compound **8** was produced from the treatment of triol **3** with 2,2-dimethoxylpropane and a catalytic amount of *p*-toluenesulfonic acid in toluene. The experimental procedures were described in Materials and Methods section.

**Scheme 1.**
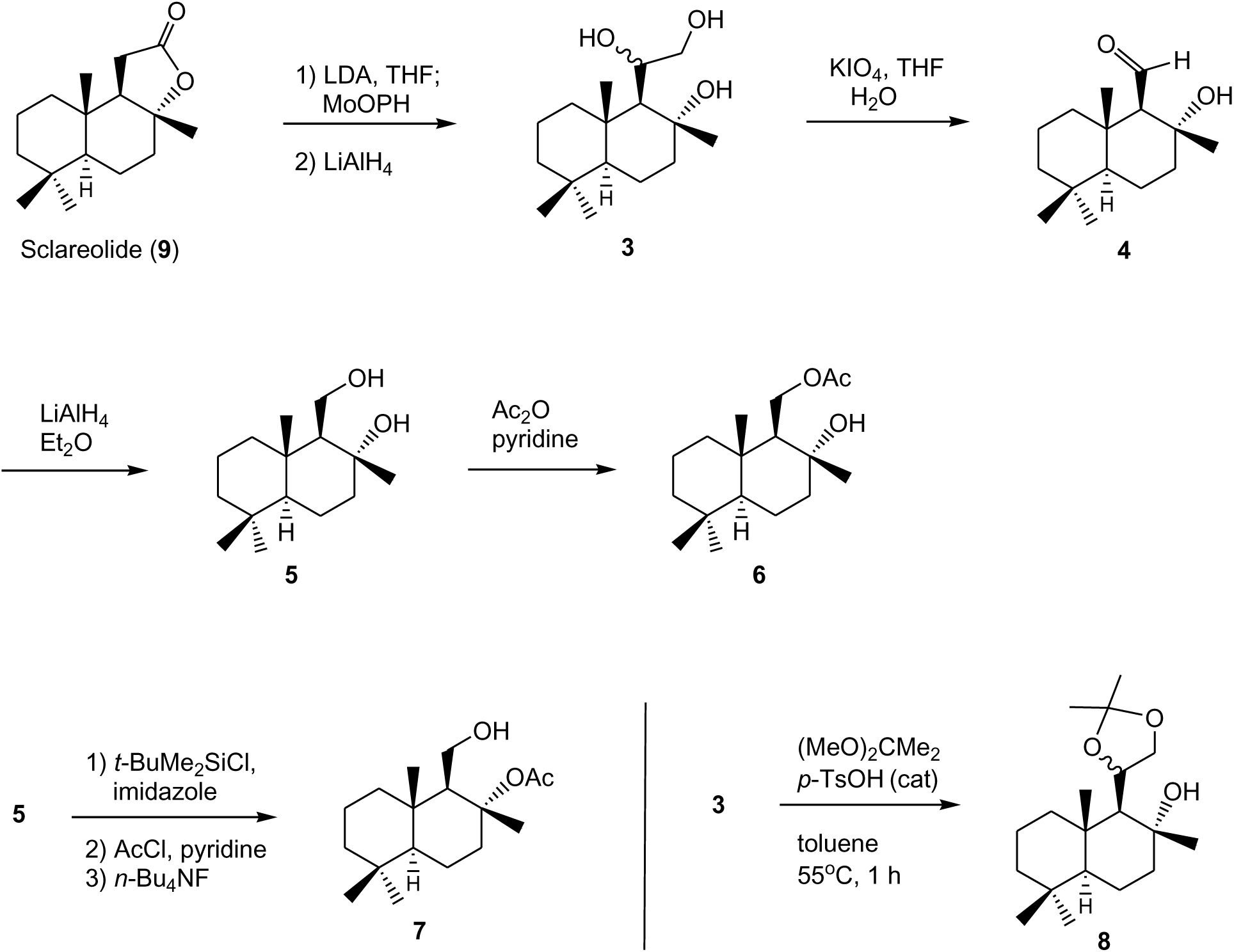
Synthesis of drimane sesquiterpenoids 3 – 8.

Examinataion of the antifungal activities of these molecules along with the mechanistic study of the most active molecule may allow future improvement in bioactivity and reduction in toxicity. We used *C. albicans* strain SC5314 for our initial screening of antifungal activities of drimane sesquiterpenoids **1** – **8**. The compounds were solubilized in dimethyl sulfoxide (DMSO), 10 mg/ml, as stock solutions and stored at −20°C. Prior to assays, stock solutions of compounds were diluted to 200 – 12.5 µg/ml in the growth media for yeast antifungal assays using the Clinical and Laboratory Standards Institute (CLSI) M38-A2 method (17). Fortuitously, we found (-)- drimenol (**1**) and (+)-albicanol (**2**) along with compound **4**, inhibit *C. albicans* SC5314 growth (Table 1). Among these three compounds, we identified **1** being more potent (with MIC value ∼30 µg/ml) than other compounds (∼60 µg/ml); therefore **1** was used for further studies including mechanistic investigation.

**Table 1.**
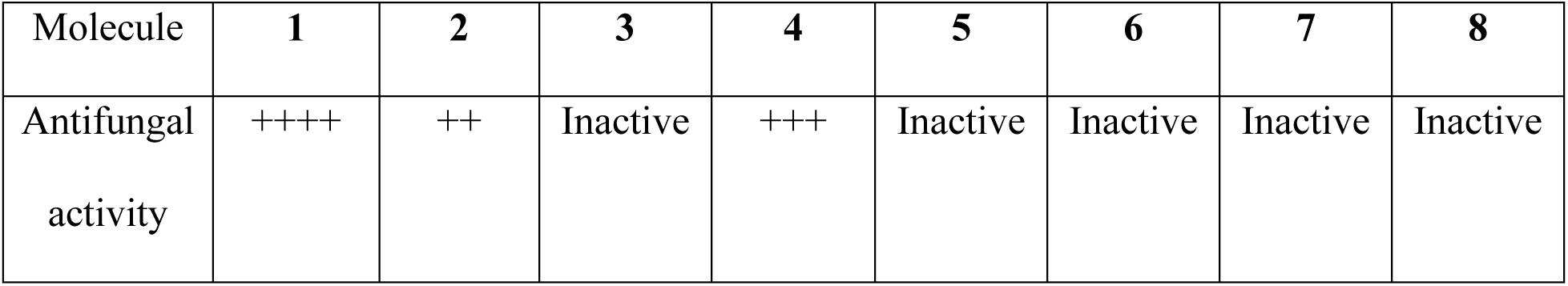
Antifungal activities of drimane sesquiterpenoids **1** – **8** against *C. albicans* SC5314. Highest activity: ++++; medium activity: +++; and low activity: ++.

| Molecule | **1** | **2** | **3** | **4** | **5** | **6** | **7** | **8** |
| --- | --- | --- | --- | --- | --- | --- | --- | --- |
| Antifungal activity | ++++ | ++ | Inactive | +++ | Inactive | Inactive | Inactive | Inactive |

### Antifungal activities of drimenol against various pathogenic fungi

Since our initial assay with *C. albicans* confirmed the antifungal activities of **1**, we extended the susceptibility assays to other pathogenic fungi including FLU resistant *C. albicans*, various species of candida, *Cryptococcus*, *Aspergillus* and a dermatophyte fungus. The CLSI broth dilution methods of M27-A3 for yeasts and M38-A2 for filamentous fungi (17) were used to determine the susceptibility. Molecule **4** was not investigated due to the presence of an aldehyde function, which may react with biological molecules.

Briefly, yeast cells or conidia (for filamentous fungi) were suspended in RPMI 1640 medium to a final concentration of 10^5^ cfu/ml and distributed in 96-well microplate to a total volume of 100 μl/well. Molecules **1** was added to the wells and two-fold serial dilutions prepared. Duplicates were used for each concentration and wells with or without DMSO served as controls. Plates were incubated without shaking at 37°C for 24 – 48 h for yeasts and 30°C for 4 days for filamentous fungi (*Aspergillus* and *Trichophyton* sp). The MIC was defined as the lowest compound concentration at which no growth occurred, as determined visually and microscopically (an inverted microscope). The results are shown in Fig. 2 and some (*C. glabrata* [BG2], *C. albicans* [FLU resistant], *C. auris, T. equinum*) in Table 2.

**FIG. 2.**
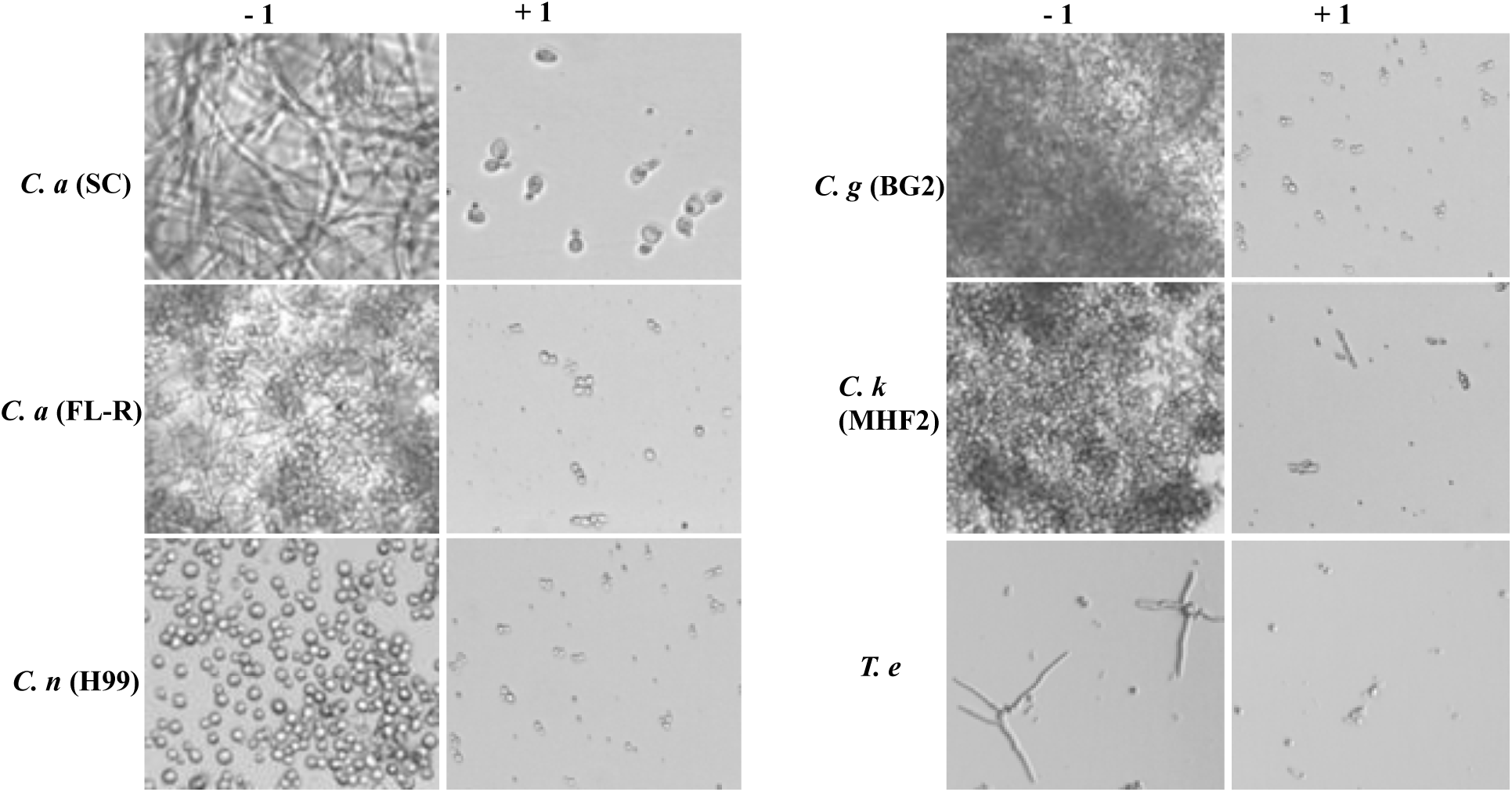
Antifungal activity of **1** against various human pathogenic fungi. The CLSI broth dilution methods of M27-A3 for yeasts and M38-A2 for filamentous fungi were used. Drimenol (**1**) was dissolved in DMSO and a two fold serial dilution was used between 200 - 12.5 μg/ml. Representative microscopic images (Leica, inverted microscope) taken at 200X magnification from **1** with cells exposed 50 μg/ml (MIC), except *T. equinum* which was exposed to 15 μg/ml at 30 °C, are shown. The MIC for **1** is 50 μg/ml under these assay conditions except mentioned otherwise. *C. a* (SC), *Candida albicans* (SC5314); *C. a* (FL-R), *C. albicans* FLU-resistant; *C. g* (BG2), *C. glabrata* (BG2); *C. k.* (MHF2), *C. krusei* (MHF2); *C. n* (H99), *Cryptococcus neoformans* (H99); *T. e, Trichophyton equinum*.

**Table 2.** Drimenol (**1**) activities against various pathogenic fungi. The MIC (100% growth inhibition) of **1** ranges from 8 μg/ml to 64 μg/ml. Fluconazole, posaconazole, and voriconazole were used as controls.

| Antifungal |  | Drimenol |  | Fluconazole | Posaconazole | Voriconazole |
| --- | --- | --- | --- | --- | --- | --- |
| Species | Isolate No. | 50% | 100% | 50% | 100% | 100% |
| *C. parapsilosis* | CLSI QC | 32 | 32 | 1 | - | - |
| *C. krusei* | CLSI QC | 32 | >64 | 16 | - | - |
| *P. variotii* | CLSI QC | 16 | 32 | - | $\leq 0.03$ | 0.125 |
| *C. albicans* | CA1 | 32 | 32 | 0.125 | - | - |
|  | CA2 | 32 | >64 | 0.25 | - | - |
|  | CA3 | 32 | 32 | >64 | - | - |
| *C. neoformans* | CN1 | 16 | 32 | 4 | - | - |
|  | CN2 | 8 | 64 | 64 | - | - |
|  | CN3 | 16 | 32 | 64 | - | - |
| *A. fumigatus* | AF1 | 16 | 32 | - | - | 0.5 |
|  | AF2 | 8 | 32 | - | - | 2 |
|  | AF3 | 16 | 32 | - | - | 4 |
| *Fusarium* | FO1 | >64 | >64 | - | - | 4 |
|  | FO2 | >64 | >64 | - | - | 8 |
|  | FS1 | 32 | 64 | - | - | 8 |
| *Scedosporium* | LP1 | 16 | >64 | - | - | >16 |
|  | SA1 | 16 | >64 | - | - | 1 |
|  | SB1 | 16 | >64 | - | - | 2 |
| *Rhizopus* | RA1 | 32 | 32 | - | 0.25 | - |
|  | RA2 | 32 | 32 | - | 0.25 | - |
|  | RA3 | 64 | >64 | - | 0.25 | - |
| *Apophysomyces* | AP01 | 32 | >64 | - | ≤0.03 | - |
|  | AP02 | 32 | >64 | - | 0.25 | - |
| *Saksenaea* | SAK1 | 16 | 16 | - | ≤0.03 | - |
|  | SAK2 | 32 | 64 | - | 0.06 | - |
|  | SAK3 | 4 | 32 | - | ≤0.03 | - |
| *Blastomyces* | BD1 | 8 | 8 | - | - | 0.5 |
|  | BD2 | 4 | 16 | - | - | 0.06 |
|  | BD3 | 4 | 8 | - | - | ≤0.03 |
| *C. glabrata* | BG2 | 30 | 50 | - | - | - |
| *C. albicans* | 95-98-flu<br>resistant | 30 | 50 | - | - | - |
| *C. auris* |  | 30 | 50 | - | - | - |
| *Trichophyton*<br>*equinum* |  | - | 15 | - | - | - |

Results summarized in Fig. 2 show that **1** has broad-spectrum fungicidal activity against various fungi including FLU resistant *C. alibcans* and *Cryptococcus* sp albeit, at a higher (50 μg/ml) MIC concentration. However, for a dermatophyte fungus, the MIC value was lower, 15 μg/ml. When compared to DMSO controls, fungi exposed to **1** showed absence of fungal growth. At higher concentration of **1** (100 μg/ml), *C. albicans* yeast cells lysed and released their cellular contents (Fig. 3, arrow). Consistent with this observation, **1** inhibited the germination of *A. nidulans* spores and appeared to cause swelling of germinating spores (Fig. 3, lower right).

**FIG. 3.**
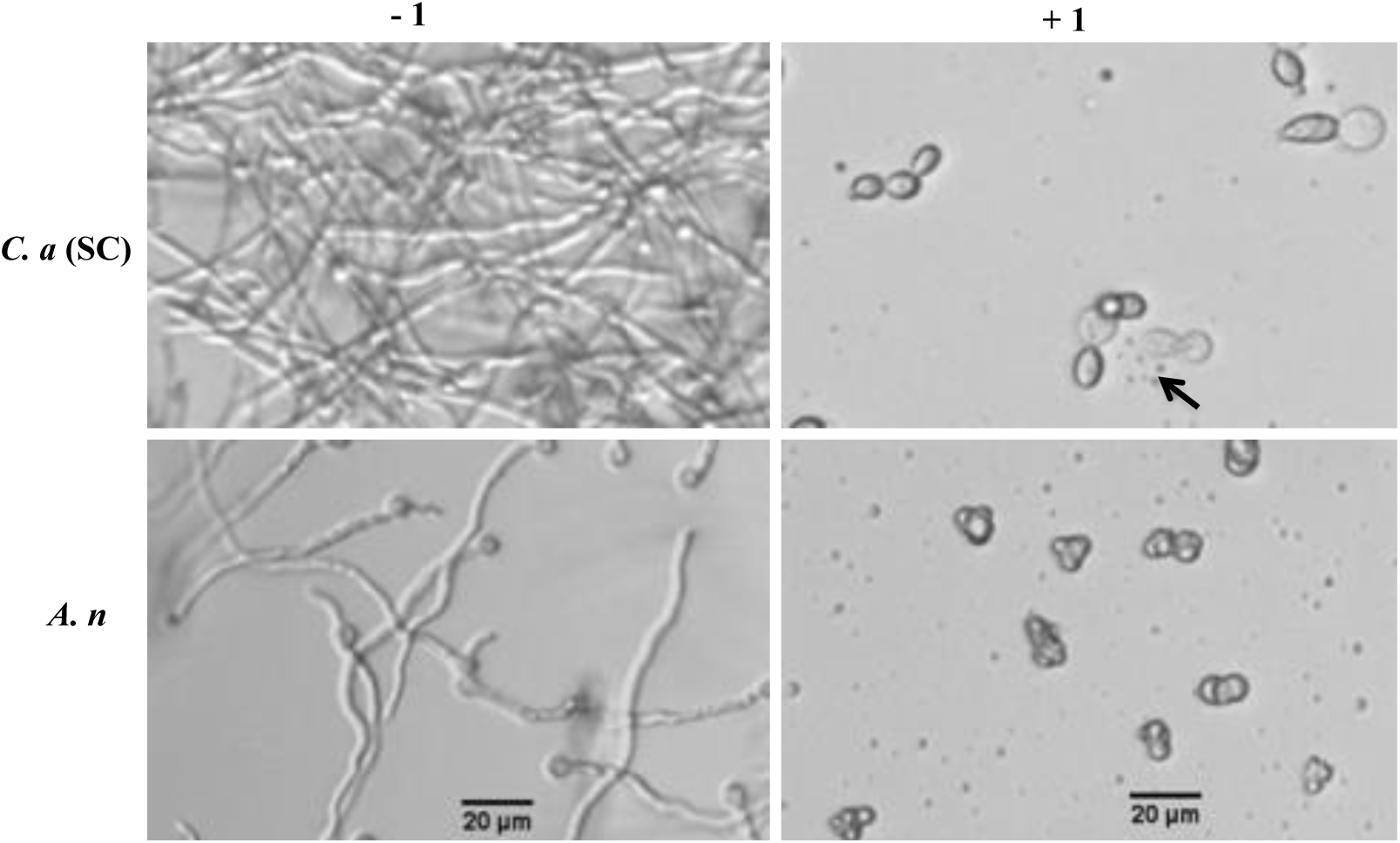
Antifungal activity of **1** against *C. albicans* and *A. nidulans*. The CLSI broth dilution methods of M27-A3 for yeasts and M38-A2 for filamentous fungi were used. *C. albicans* (SC5314) showed lysis of yeast cells (arrow) at 100 μg/ml of **1** compared to the control where a network of hyphal growth was observed. Similarly, the germination of *Aspergillus nidulans* spores was inhibited by **1** (lower, right panel). Representative microscopic images (Leica, inverted microscope) were shown. Scale bar, 20 μm.

To extend our antifungal screening with **1** against additional human pathogenic fungi, we have used the non-clinical and pre-clinical service program offered by the NIH NIAID supported fungus testing center at the University of Texas Health Sciences Center, San Antonio. The fungus testing center used the CLSI M38-A2 method (17) to determine the MIC of **1** after 24 - 72 hours incubation with concentrations ranging from 0.125 - 64 μg/ml. Positive control antifungals (FLU, posaconazole and voriconazole) were also included in parallel. The results are summarized in Table 2.

Next, we determined the viability of fungal cells that were exposed to **1**. Cells exposed to 50 μg/ml (MIC) for 24 h or at 100 μg/ml for 48h were used. To determine the viability of treated fungal cells, small volumes (1 - 5 μg/ml) of mixed cell suspensions were removed from wells and spotted on YPD agar medium. The agar plates were incubated at 30°C for 24 h - 72 h and the growth of fungi were recorded. Growth of yeasts occur in 24 h and filamentous fungi in 48 - 72 h for control (without **1**) but not for those treated with **1** suggesting that it acts as a fungicidal compound (data not shown).

### Drimenol acts better than fluconazole against *Candida auris* growth

*C. auris* is an emerging multidrug resistant fungal pathogen that is known to cause nosocomial infections with ‘superbug”-like traits (18). *C. auris* was first discovered in 2009 in Southeast Asia and now it is present in 33 countries across 6 continents. Since this fungus is resistant to all antifungals and is invasive, the mortality rate is high (19). Recently, CDC has issued a clinical health emergency warning about this fungus. Since **1** showed a broad-spectrum fungicidal activity, we determined its effect against *C. auris* growth using a bioscreen-C growth monitoring system. *C. auris* was grown in the presence or absence of **1** in RPMI medium (CLSI method) (17) for 24 h at 37° C. Negative controls (solvent) and positive controls (FLU) were included in parallel. Results depicted in Fig. 4 indicate that **1** inhibited *C. auris* growth completely at 60 μg/ml. In contrast, FLU at the same concentration (60 μg/ml) showed poor inhibition of growth. Thus, **1** could be useful as a broad-spectrum fungicidal compound.

**FIG. 4.**
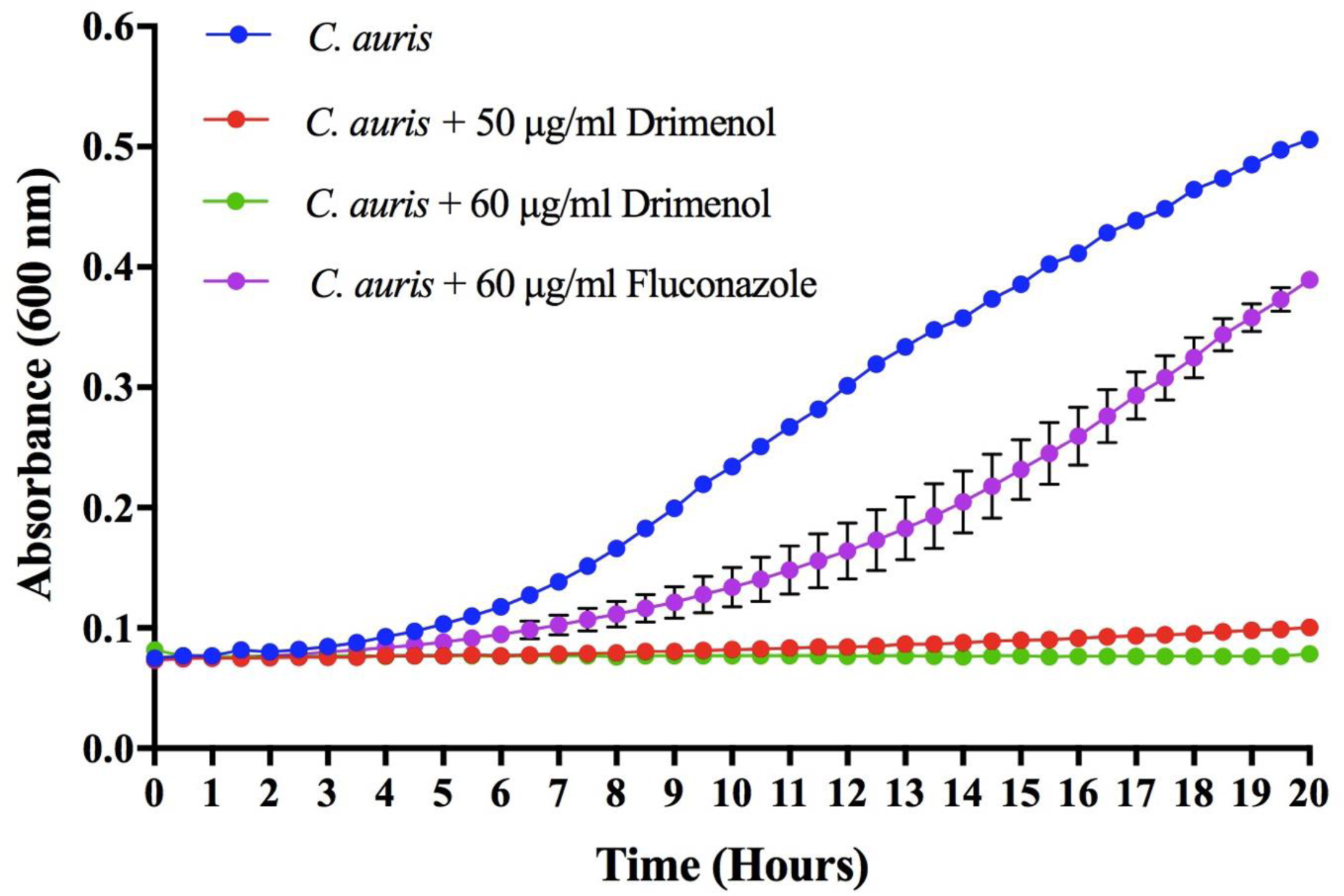
**Drimenol** (**1**) **inhibits *C. auris* growth better than fluconazole**. *C. auris* was grown in honeycomb microtiter wells containing RPMI medium in the presence and absence of **1** for 20 h at 37° C. Fungus growth was measured by absorbance at OD 600 nm using Bioscreen-C growth monitor. Growth curves showed the mean of triplicates and experiments were repeated at least twice. Error bars are SD and were too short to appear in the line graphs.

### Drimenol (1) is tolerated by *C. elegans* and protects it from fungal mediated death

Invertebrate animal models provide an inexpensive and powerful platform to test antifungal compounds for their efficacy and toxicity simultaneously. We evaluated **1** for its antifungal activity and tolerance in *C. elegans* infection model of candidiasis as described before (20). Results shown in Fig. 5 indicate that **1** can protect worms from *C. albicans* mediated death and that the worms were not adversely affected by **1**, as judged by their motility and viability following compound exposure.

**FIG. 5.**
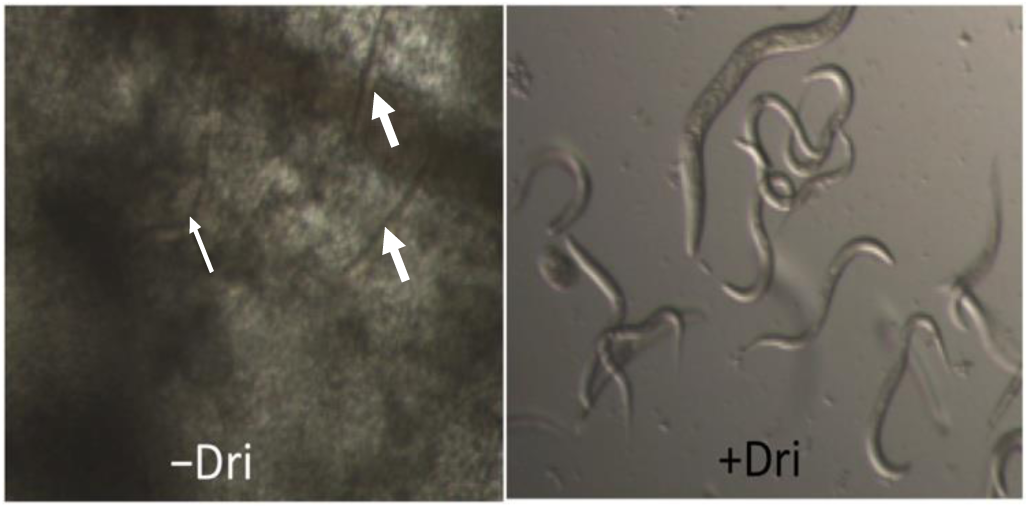
**Protection of *Caenorhabditis elegans* worms from *C. albicans* mediated death by drimenol**. *C. albicans* (yeast cells) fed larvae were incubated in RPMI medium without and with **1** (+Dri) (50 µg/ml) in a 96 well microtiter plate and incubated at 30 °C for 2 – 3 days. Left panel without drimenol shows died worms (straight and immobile, thick arrows) due to *C. albicans* growth. Thin arrow shows a weakly moving worm. Right panel shows **1** containing well where worms are alive as judged by their movements and the lack of fungal growth.

### Mechanism of drimenol (1) antifungal activity

To understand the compound’s mechanisms of action (MOA), researchers have used pooled library of genome-wide barcoded mutant collections of *Saccharomyces cerevisiae* or *C. albicans* for drug-induced sensitivity assay or the haploinsufficiency (HIP) assay (21–23). For example, if **1** can inactivate partially or completely its protein target in the heterozygous mutant pool, the resulting growth defect of that mutant(s) can be measured quantitatively by sequencing the tagged unique barcodes. This approach will help narrowing down the putative target(s). Similarly, a homozygous nonessential mutant library can be used as a complementary approach to the heterozygous essential mutant collection to verify the target pathway/genes of compounds. In this case, if the homozygous mutant of a gene is sensitive to the compound, then that gene may not be the drug target (21) as the homozygous mutant lacks the gene product. This imples that the compound may exert its effect via drug-induced synthetic lethality. Thus, by combining data from both heterozygote and homozygote screens one may determine the compound’s MOA.

In this study, we used *S. cerevisiae* barcoded homozygous nonessential and heterozygous essential, and *C. albicans* barcoded heterozygous Tn mutant (23) libraries. Briefly, IC-50 of **1** for *C. albicans* and *S. cerevisiae* was determined in yeast growth (YPD) condition (Fig. 6). Based on this assay results, IC-50 of 25 μg/ml for *C. albicans* and 15 μg/ml for *S. cerevisiae* was calculated for **1**. Two different sub-MIC concentrations of **1** were selected for determining the mechanism of action against mutant libraries.

**FIG. 6.**
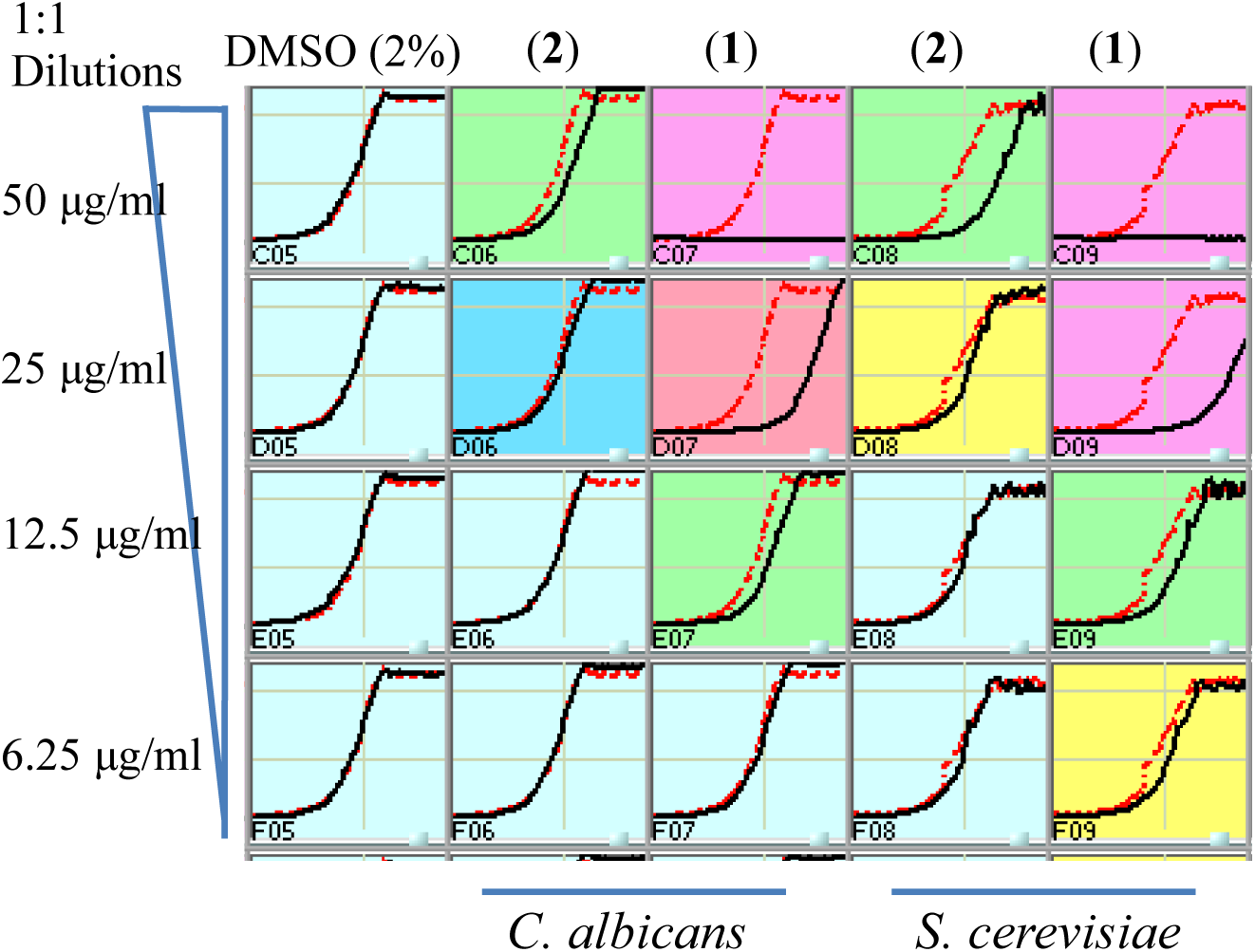
**Determination of IC-50 for drimenol** (**1**) **activity against *C. albicans* and *S. cerevisiae***. Yeast growth conditions (YPD medium at 30°C) were used to determine the IC-50 values. The red line in each panel indicates growth of the DMSO reference. An IC-50 of ∼25 μg/ml for *C. albicans* and ∼15 μg/ml for *S. cerevisiae* were calculated for **1**. Albicanol (**2**) showed weaker activity against both fungi and was not considered for further analysis.

Next, pooled *S. cerevisiae* and *C. albicans* mutant collections were grown separately in the presence or absence of compounds (with DMSO) for 20-generations, barcodes from genomic DNA were amplified and relative strain abundance were quantified based on TAG microarray signals. The log_2_ ratio of tag signals between DMSO control and **1** exposed samples were presented in scatter plots as the “fitness defect” (Figs. 7A & B). Mutants that were depleted from the growth pool due to **1** are indicated by circles. Mutants that were highly susceptible to **1** are shown with high log ratio (*e.g*. SEC66 in Fig. 7A) (highly depleted in the pool) and considered putative targets. Lists of *S. cerevisiae* and *C. albicans* mutants that are highly sensitive to **1** were given in **Supplementary Tables 1 and 2,** respectively.

**FIG. 7.**
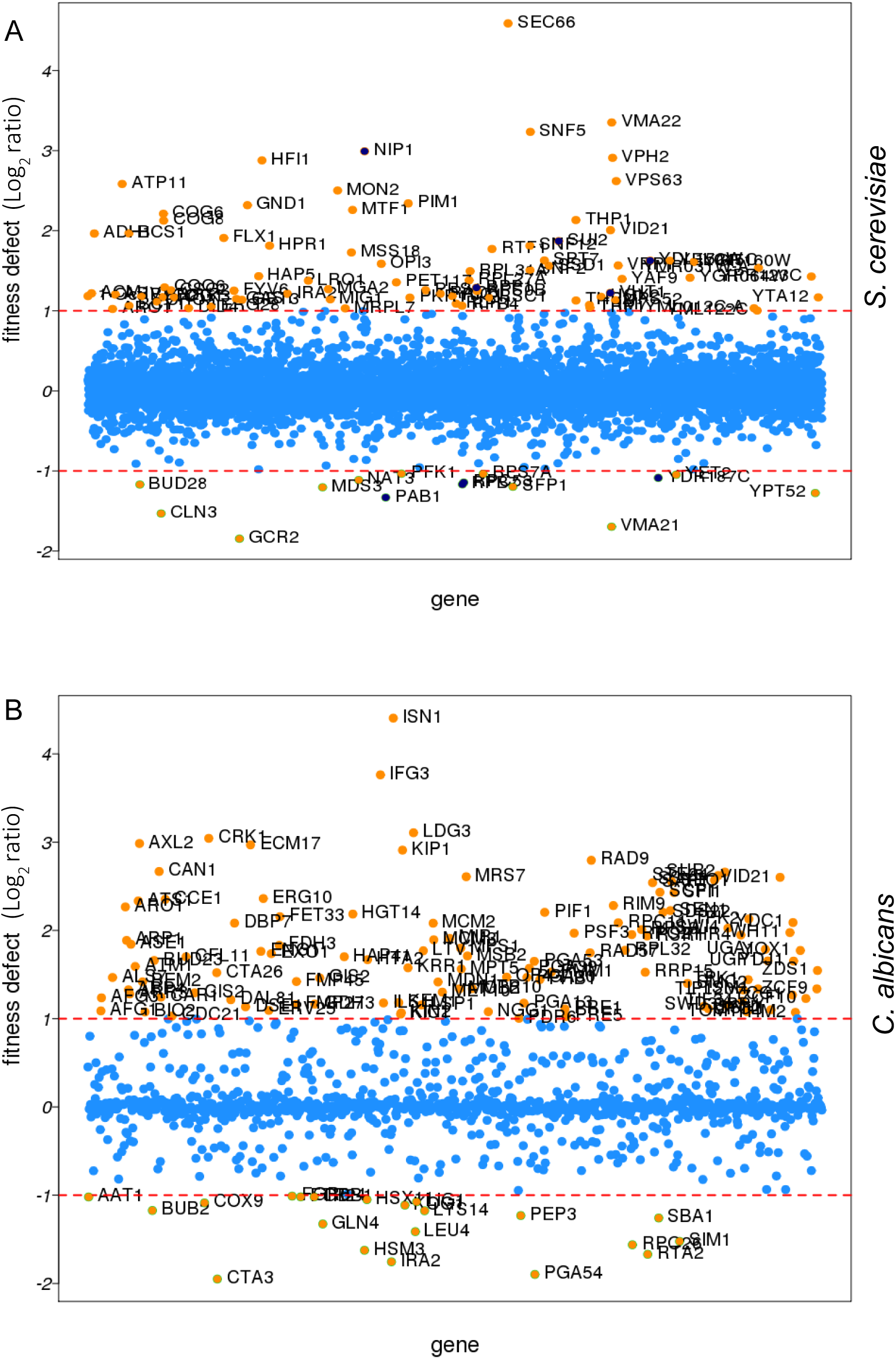
**Genome-wide screens of *S. serevisiae* (A) and *C. albicans* (B) mutant libraries against drimenol (1) for drug induced hypersensitivity.** Pooled collections of *S. cerevisiae* nonessential homozygous and essential heterozygous mutants were grown in the presence and absence of **1** at the concentration of 0.025 mg/ml for the indicated number of generations before profiling for their abundance (DNA barcodes). Twenty generations for essential heterozygous and five generations for non essential homozygous mutants were used which gave an optimum of ∼20% growth inhibition. Similarly, *C. albicans* Tn-insertion mutants (heterozygous, 20 generations) was used at 0.025 mg/ml of **1.** Each spot represents single mutant. The log ratio of each mutant (**1** exposed vs no drug control) was calculated and presented in scatter plots where greater the number the more sensitive that strain is.

Our results of forward genetic screening from mutant libraries of *C. albicans* and *S. cerevisiae* with **1** indicate that it affects cellular activities involved in protein secretion, vacuolar functions, chromation remodeling and cyclin dependent protein kinases (CDK) (Fig. 7 and **Supplementary Tables 1 & 2**). For example, SEC66, highly sensitive to **1** (log_2_ >4.5, Fig. 7A), is a component of Sec63 SECretary complex in *S. cerevisiae* involved in protein targeting and importing into the ER. Similarly, VMA22 is a vacuolar membrane ATPase required for vacuolar H+-ATPase function and localized to the yeast ER (Saccharomyces Genome Database).

The *ISN1* (log_2_ 4.4, Fig. 7B) gene product is involved in inosine 5’-monophosphate 5’-nucleotidase activity in *C. albicans* (Candida Genome Database). This gene product is uncharacterized and it is present only in fungi and not in human or murine, suggesting that Isn1p a suitable antifungal drug target. *IFG3* is a putative D-amino acid oxidase, which is uncharacterized, and *CRK1* is a protein kinase of the Cdc2 subfamily involved in hyphal development and virulence in *C. albicans* (24). The *CRK1* ortholog in *S. cerevisiae* is *SGV1,* which is a part of BUR2 kinase complex and plays a major role in transcriptional regulation.

### Yeasts spot assay to validate drimenol (1) mechanism of action

Based on the forward genetic library screening assay results (Fig. 7) and their functions inferred from the available literature, we selected few heterozygous mutants of *C. albicans* that had high to medium positive log_2_ ratio (hypersensitive, *CRK1* and its putative interacting partners proteins

*CDC37, Orf19.759, Orf19.1672 and Orf19.4382*) and few with negative log_2_ ratio (resistance, *VPS53*, *TSC11* & *PHO89*) to verify the genetic screening data. Agar medium containing sub-MIC concentration of **1** (30 μg/ml) or solvent was used to spot test suspensions of various mutants (GRACE mutant collection) (25).

Results shown in Fig. 8 confirm the findings of forward genetic screening assays of *C. albicans* mutants. For example, Crk1 Kinase is predicted to associate with *RET2*, which involves in retrogradetion of vesicle transport for protein signaling and secretion (*SEC*66), and plays a role in protein translation with G1/S cell-cycle transition (26). Products encoded by Ret2, orf19.759 (*SEC21*), orf19.1672 (*COP*1) and orf19.4382 (*RET3*) are uncharacterized and are likely targets of Crk1 kinase, which are defective in growth on agar medium containing **1** (Fig. 8). Molecule **1** induced hypersensitivity of these mutants, which represent candidate targets. The orf19.759 (*SEC21* ortholog of *S. cerevisiae*) is uncharacterized in *C. albicans*. Sec21 involves in transport from endoplasmic reticulum (ER) to Golgi vsicle-mediated transport (anterograde), Golgi to ER (retrograde) and COPI vesicle coat, and endosome localization (27). *RET2* is also uncharacterized in *C. albicans* and the ortholog is a subunit of the coatomer complex (COPI), which coats Golgi-derived transport vesicles, involves in retrograde transport between Golgi and ER, and interacts with Crk1 kinase in the two-hybrid system (28). Crk1 is known to play a role in regulating trafficking and secretion of effectors by interacting with the early endosome during *Ustilago maydis* (corn smut fungus) infection in corn plants (29, 30).

**FIG. 8.**
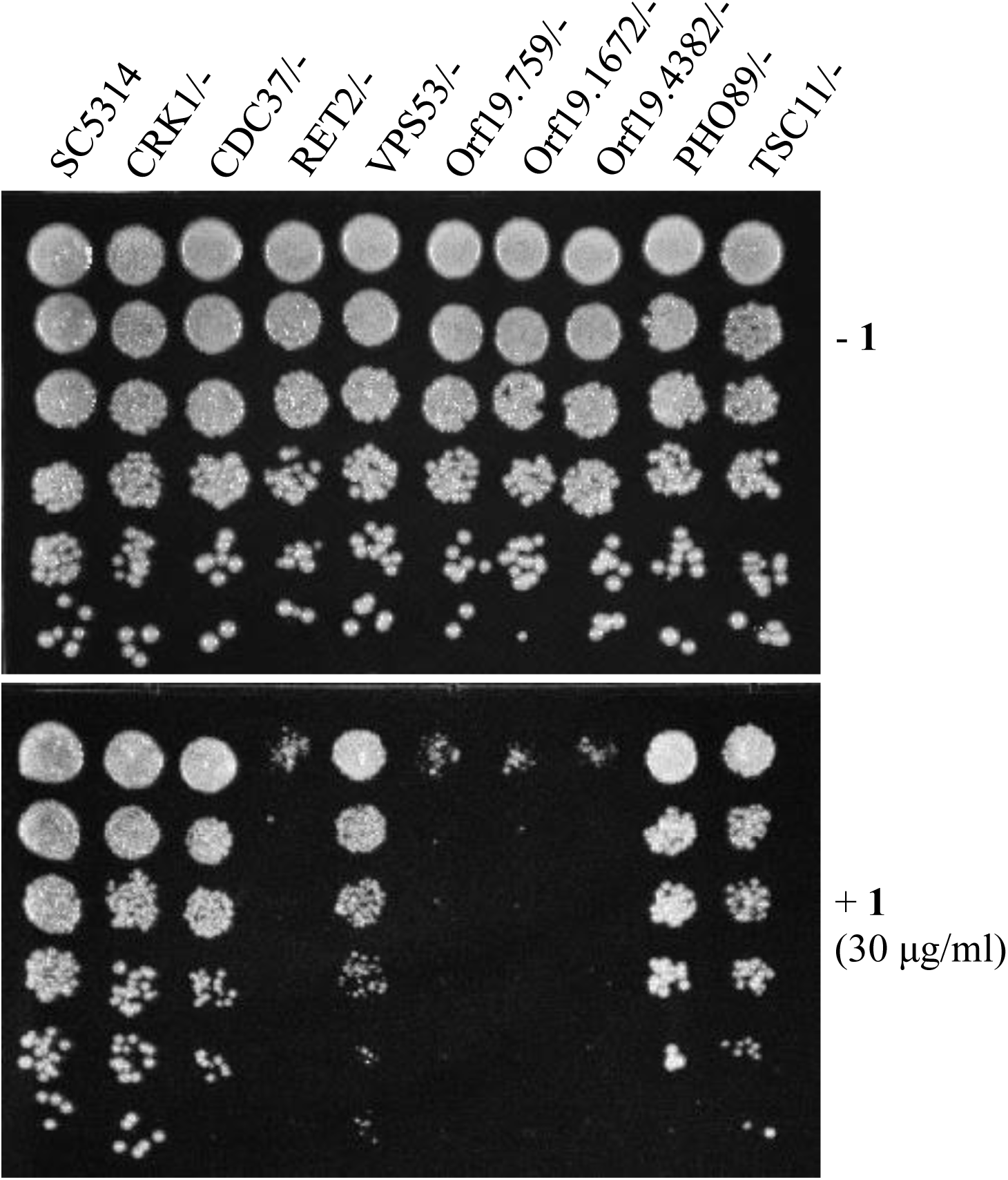
**Validation of *C. albicans* Tn-insertion mutant screen data by yeast spot assay**. A four-fold serially diluted yeast cultures of indicated *C. albicans* heterozygous mutants (GRACE) were spot tested on YPD agar containing **1** (30 μg/ml) or DMSO (**-1**). Heterozygous mutants (*RET2/-, Orf19.759/-, Orf19.1672/-*, and *Orf19.4382/-*) affected by **1** directly or indirectly were hypersensitive and showed lack of growth.

Since Crk1 kinase may interact with multiple targets (Ret2, orf19.759, orf19.1672 and orf19.4382) and because Crk1 represents an important antifungal drug target (24), we performed computational molecular docking of **1** with *Cryptosporidium parvum* Crk1 crystal structure (2QKR-A) (31) using AutoDock Vina software (32). Results showed in Fig. 9, suggest that **1** can interact with the *N*-terminal catalytic domain of *C. albicans* Crk1 (which has 61% similarity and 40% identical to the *C. parvum* Crk1). Particularly, noteworthy from our computational docking studies is that **1** has close interactions with Gly 31, Val 37, Gln 148, Leu 151, and Phe 98 amino acid residues. The docked structure shows that an available open space in Crk1 for incorporation of an additional function group onto the cyclohexane A ring of **1** (Fig. 1), signifying a possible modification of **1** for future improvement of biological activity. Thus, this CDK member may comprise the target of **1** and it is notable that this conserved gene is present in many of the tested pathogenic fungi (Table 2).

**FIG. 9.**
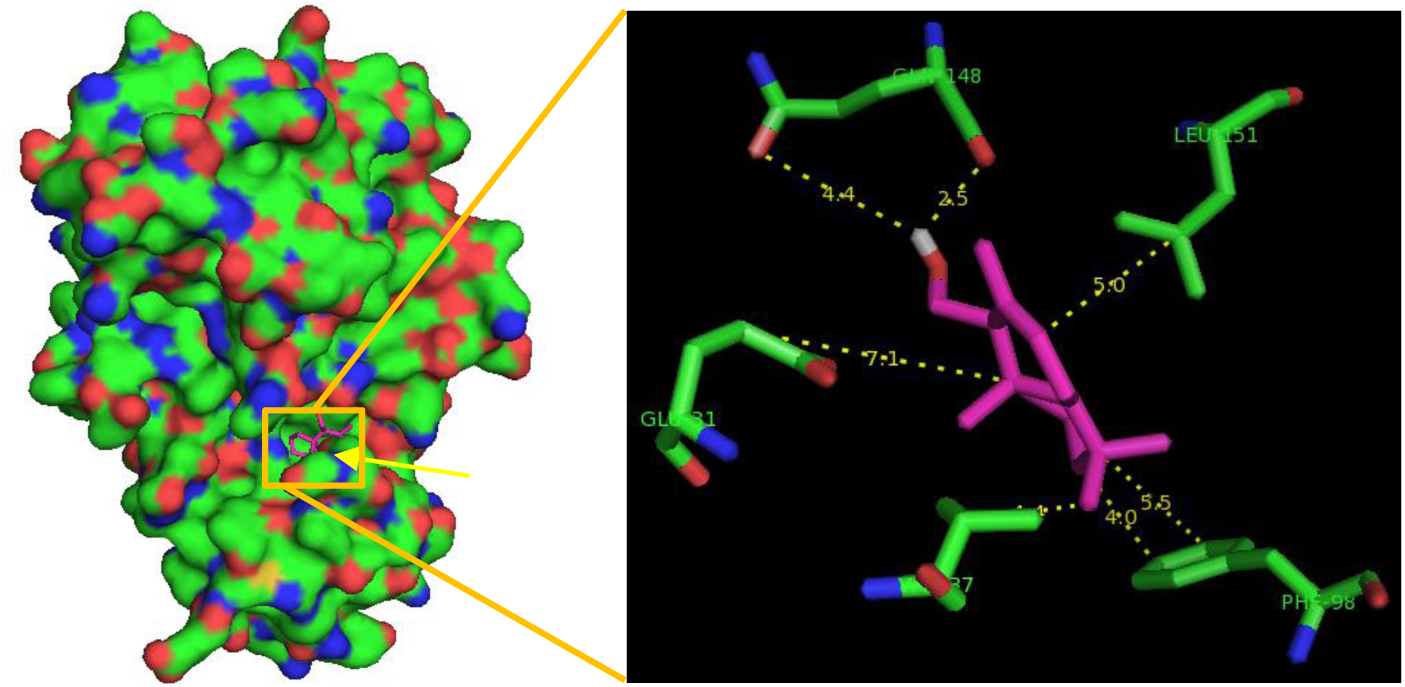
**Molecular docking of drimenol with *C. parvum* Crk1 kinase.**

## DISCUSSION

Fungi have emerged in the last two decades as major causes of human disease. *C. albicans* is a major fungal pathogen affecting at all ages and fourth leading cause of nosocomial bloodstream infections in the US (33). *C. albicans* and other *Candida* spp cause mucosal, disseminated and invasive candidiasis, especially among patients who are immunocompromised or hospitalized with serious underlying diseases. The overall mortality for invasive diseases caused by *Candida* spp. and *Aspergillus* spp. is around 50% (33, 34). While there are more than 150 species of Candida, about 15 species are recognized as frequent human pathogens (34, 35). Some of them are: *C. albicans, C. glabrata, C. krusei, C. tropicalis* and *C. parapsilosis*. Among these, *C. albicans* is by far the most common species isolated from humans and is a frequent denizen of the oropharynx, mucousal surfaces, gastrointestinal and genitourinary tracts. In the developing world, there are ∼1 million cases of cryptococcal diseases per year resulting in 675,000 deaths (9, 36). *Cryoptococcus neoformans* is an opportunistic fungal pathogen that causes meningitis in immunocompromised individuals. Often found in soils contaminated with bird feces, *C. neoformans* enters its host through the lungs via inhalation of spores. Some of the cryptococcal species are hypervirulent (37) and have drawn a considerable public attention due to their causative role in the cryptococcosis outbreak throughout the Pacific Northwest (38, 39). Only few antifungals can be useful to treat cryptococcosis and drug resistant strains are emerging.

*Aspergillus* spp are ubiquitous molds found widely in the environment as saprophytes and produce microscopic spores or conidia which upon inhalation, cause invasive pulmonary disease. In immunocompromised patients such as hematopoietic stem cell transplantation, solid organ transplantation and undergoing chemotherapeutic agents or immunomodulatory agents, invasive aspergillosis remains the most important cause of infection-related mortality (40, 41). Among several species of Aspergillus, *A. fumigatus* and *A. flavus* are frequent pathogens. Dermatophytes are another group of keratinophilc pathogenic fungi that causes variety of infections in humans and animals (42). Some of these fungi include *Trichophyton tonsurans* (scalp ring-worm), *T. equinum*, and *Microsporum gypseum* (garderner’s ringworm). Emerging fungal disease such as zygomycosis is life-threatening particularly during natural calamity (*e.g.* the 2004 tsunami, the 2008 Katrina and May-2011 Joplin tornado). Novel compounds with broad-spectrum antifungal activity are highly desirable to treat various fungal pathogens.

Because fungi are eukaryotes, the development of antifungal therapeutics that are nontoxic to human is challenging due to the availability of relatively few targets. In the last thirty years, only one new class of antimycotic (β-glucan synthase inhibitor, the echinocandins) was introduced into clinical practice. Although this drug is an important addition, it has a number of limitations including ineffectiveness against *Cryptococcus sp* and poor oral bioavailability (43). New drugs are needed to be discovered and developed.

In our search for novel antifungal small molecules from our available synthetic terpenoids, we have identified two compounds, (-)-drimenol (**1**) and (+)-albicanol (**2**) (Fig. 1), that showed strong activity against *C. albicans*. Among these two compounds, **1** showed stronger bioactivity. It acts not only against *C. albicans* as fungicidal but also against *Aspergillus nidulans*, FLU resistant strains of *C. albicans*, *C. glabrata, C. krusei*, *Cryptococcus* spp and other dermatophytes, suggesting that **1** is a broad-spectrum antifungal agent (Figs. 2, 3 and Table 2). At an increased concentration (100 μg/ml), **1** caused rupturing of the fungal cell wall/membrane, *e.g*. *C. albicans* (Fig. 3) and *Cryptococcus sp.* (data not included). *Candida auris* is an emerging and multi antifungal resistant strain that causes nosocomial infection and has been reported recently across the world (18). Our bioscreen-based growth curve monitoring assay with **1** showed better activity than clinical antifungal drug FLU (Fig. 4) indicating a potential use of **1** against *C. auris* and other drug-resistant fungal pathogens.

Molecule **1** is a natural product presents in liverworts and higher plants (44, 45), and its antifungal mechanism against human pathogenic fungi has not been reported previously. A recent study has shown that **1** has antifungal activity against *Botrytis cinerea*, a plant fungal pathogen and the mechanisms appear to act by fungal membrane damage and reactive oxygen species (ROS) production (46).

In order to develop broad-spectrum novel antifungal compounds, we evaluated **1** against various fungi that are pathogenic to humans and determined its mechanisms of action in *C. albicans* and *S. cerevisiae*. Based on our yeasts mutant screening data and subsequent spot assay results, we found that **1** acts as a fungicidal compound by affecting cellular activities targeting protein trafficking between Golgi to ER, protein secretion (Sec system) and cell signaling, possibly through cell division related kinase 1, Crk1 (Figs. 7 & 8). Genetic methods have been used to determine the mechanism of antifungal compounds by drug-induced hypersensitivity assay (22, 47). Using similar approaches, we showed that **1**-mediated inhibition of *C. albicans* heterozygous mutants of *CDC37, Orf19.759, Orf19.1672 and Orf19.4382*, the known or putative targets of Crk1 kinase, at sub-MIC concentration. In support of this observation, computational molecular docking of **1** with the crystal structure of a fungal (*C. parvum*) Crk1 kinase showed interactions of **1** with the key residues in the catalytic domain (*N*-terminal) of Crk1 (Fig. 9).

Cinnamodial is a closely related compound belong to the drimane sesquiterpenoid family with potent antifungal activity (48), but its chemical structure (containing dialdehyde groups; Fig. 1) and physiological properties are quite different from **1**. For example, the antifungal activity of cinnamodial was shown to abolish by amine compounds (likely due to a coupling reaction from the aldehyde functions of cinnamodial with the amino group of amine compounds) or when cinnamodial was incubated in YPD medium (19, 49). In contrast, molecule **1**’s bioactivity was not affected by amines or YPD medium (Fig. 5). Thus, the antifungal mechanisms of **1** could be different from cinnamodial. Since the synthetic route for **1** and its analogs are well established, improvements of its antifungal properties are possible through medicinal chemistry approaches. In summary, we have synthesized a focused library of drimane sesquiterpenoid compounds and identified **1** as a broad-spectrum fungicidal compound against various human pathogenic fungi including *C. albicans, C. auris, Cryp. neoformans*, *Aspergillus, Blastomyces, Scedosporium, Fusarium, Pneumocystis*, and dermatophytes at 8 - 64 μg/ml. By employing the libraries of bar-coded *C. albicans* and *S. cerevisiae* genome-wide mutants, the mechanism of action of **1** was determined. Further evaluation of **1** in animal models of fungal diseases would help develop **1** as an antifungal agent.

## MATERIALS AND METHODS

### Synthesis of drimane molecules. (1*R*,2*R*,4a*S*,8a*S*)-2-Hydroxy-2,5,5,8a-tetramethyl-decahydronaphthalene-1-carbaldehyde (4)

To a solution of 0.20 g (0.74 mmol) of triol **3** (13) in 10 ml of THF and 2.5 ml of water was added 0.19 g (0.81 mmol) of potassium periodate. The resulting mixture was stirred at 25°C for 4 hours, diluted with water (50 ml) and extracted three times with ethyl acetate (50 ml each). The combined extract was washed with water and brine, dried (anhydrous Na_2_SO_4_), concentrated, and column chromatographed on silica gel using a mixture of hexane and ethyl acetate (20:1) as an eluent to give 0.16 g (91% yield) of compound **4**, whose spectral data is in agreement with that reported (13).

### (1*S*,2*R*,4a*S*,8a*S*)-1-(Hydroxymethyl)-2,5,5,8a-tetramethyl-decahydronaphthalen-2-ol (5)

To a cold (0°C) solution of 1.0 g (4.2 mmol) of aldehyde **4** in 80 mL of diethyl ether under argon, 80 mg (2.1 mmol) of lithium aluminum hydride was added in portions. The resulting solution was stirred at 0°C for 30 minutes, diluted with aqueous NH_4_Cl, and extracted with diethyl ether three times (50 ml each). The combined extract was washed with water and brine, dried (MgSO_4_), and concentrated to give 0.98 g (97% yield) of diol **5**, whose spectral data are in agreement with that reported (13).

### [(1*S*,2*R*,4a*S*,8a*S*)-2-Hydroxy-2,5,5,8a-tetramethyl-decahydronaphthalen-1-yl]methyl acetate (6)

To a cold (0°C) solution of 0.10 g (0.40 mmol) of diol **5** in 2 ml of dichloromethane and 0.32 g (4.0 mmol) of pyridine under argon, was added 49 μl (0.48 mmol) of acetic anhydride, and the resulting solution was stirred at 0°C for 30 minutes and 25°C for 1 h. It was diluted with 30 ml of aqueous NH_4_OH, extracted twice with diethyl ether (30 ml each), and the combined extracts were washed with water and brine, dried (anhydrous Na_2_SO_4_), concentrated, and column chromatographed on silica gel using a gradient mixture of hexane and diethyl ether as eluents to give 90 mg (80% yield) of acetate **6**. Mp. 64 – 67 °C; [α]^D_22_^ = −8.2 (c = 0.55, CHCl_3_); ^1^H NMR (CDCl_3_; 400 MHz) δ 4.35 (dd, *J* = 12, 4 Hz, 1 H), 4.24 (dd, *J* = 12, 4 Hz, 1 H), 2.05 (s, 3 H), 1.88 (dt, *J* = 12, 2 Hz, 1 H), 1.70 – 0.93 (a series of m, 11 H), 1.17 (s, 3 H), 0.88 (s, 3 H), 0.86 (s, 3 H), 0.80 (s, 3 H) ppm; ^13^C NMR (CDCl_3_; 100 MHz) δ 171.4, 72.6, 62.6, 60.0, 55.7, 44.0, 41.7, 39.7, 38.1, 33.5, 33.2, 24.6, 21.6, 21.3, 20.3, 18.4, 15.8 ppm. MS (electrospray ionization), m/z 283.1 (M+H^+^). HRMS-ESI: m/z [M + H]^+^ calcd for C_17_H_31_O_3_^+^: 283.2268, found: 283.2273.

### (1*S*,2*R*,4a*S*,8a*S*)-1-(Hydroxymethyl)-2,5,5,8a-tetramethyl-decahydronaphthalen-2-yl acetate (7)

Compound **7** was prepared by a sequence of three reactions: (i) silylation of the primary alcohol function of **5** with *t*-butyldimethylsilyl chloride; (ii) acetylation of the tertiary alcohol function with acetyl chloride and pyridine; and (iii) removal of the *t*-butyldimethylsilyl ether protecting group with tetra-*n*-butylammonium fluoride in THF.

To a solution of 9.5 mg (40 μmol) of compound **5**, 11 mg (150 μmol) of imidazole, and 6 mg (49 μmol) of 4-(dimethylamino)pyridine in 2 ml of dichloromethane under argon at 25°C, was added 14.3 mg (95 μmol) of *t*-butyldimethylsilyl chloride, and the solution was stirred for 4 h. The reaction mixture was diluted with 10 ml of aqueous ammonium chloride and extracted with diethyl ether three times (10 ml each). The combined extracts were washed with water (10 ml) and brine (10 ml), dried (anhydrous Na_2_SO_4_), concentrated to give 12.5 mg of the mono-silylated product. This crude product was used in the subsequent step without purification. To a solution of the above mono-silylated product and 0.1 ml of pyridine in 0.5 ml of dichloromethane under argon at 0°C, was added 10 μl (0.13 mmol) of acetyl chloride. The reaction mixture was stirred at 25°C for 2 h, diluted with aqueous ammonium chloride (10 ml), and extracted three times with diethyl ether (10 ml each). The combined extracts were washed with brine, dried (anhydrous Na_2_SO_4_), concentrated to give the crude product, which was used in the following step without purification. The above crude product was dissolved in 1 ml of dried THF (distilled over sodium/benzophenone) and 0.3 ml (0.3 mmol) of tetra-*n*-butylammonium fluoride (1 M solution in THF) and stirred at 25°C under argon for 1 h. The reaction solution was diluted with 0.1 N ammonium hydroxide (10 ml) and extracted with diethyl ether three times (10 ml each). The combined extracts were washed with water (10 ml) and brine (10 ml), dried (anhydrous Na_2_SO_4_), concentrated, and column chromatographed on silica gel using a gradient mixture of hexane and diethyl ether to give 4.2 mg (38% overall yield from diol **5**) of compound **7**. Compound **7**: Mp. 101 – 103 °C; [α]^D_22_^ = +0.35 (c = 0.23, CHCl_3_); ^1^H NMR (CDCl_3_; 400 MHz) δ 3.91 (dd, *J* = 12, 2 Hz, 1 H), 3.84 (dd, *J* = 12, 2 Hz, 1 H), 2.95 – 2.90 (m, 1 H), 1.98 (s, 3 H), 1.88 (dt, *J* = 12, 2 Hz, 1 H), 1.62 (s, 3 H), 1.70 – 0.88 (a series of m, 10 H), 0.94 (s, 3 H), 0.87 (s, 3 H), 0.82 (s, 3 H) ppm; ^13^C NMR (CDCl_3_; 100 MHz) δ 169.9, 84.9, 63.8, 59.8, 55.8, 41.8, 39.5, 38.2, 36.1, 33.5, 33.2, 25.8, 22.8, 21.7, 18.3 (2 C), 16.1 ppm. MS (electrospray ionization), m/z 305.1 (M+Na^+^). HRMS-ESI: m/z [M + Na]^+^ calcd for C_17_H_30_NaO_3_^+^: 305.2087, found: 305.2082.

### (1*S*,2*R*,4a*S*,8a*S*)-1-(2,2-Dimethyl-1,3-dioxolan-4-yl)-2,5,5,8a-tetramethyl-decahydronaphthalen-2-ol (8)

A solution of 18 mg (67 μmol) of triol **3**, 50 μl of 2,2-dimethoxypropane and 3 mg of anhydrous *p*-toluenesulfonic acid in 1 ml of toluene was stirred under argon at 55°C for 1 h. The solution was cooled to room temperature, neutralized with sodium bicarbonate (∼3 mg), diluted with 10 ml of water, and extracted with ethyl acetate three times (15 ml each). The combined extract was washed with brine, dried (MgSO_4_), concentrated and column chromatographed on silica gel using a gradient mixture of hexane and diethyl ether as eluent to give 14 mg (71% yield) of compound **8** as a mixture of two stereoisomers: (the major isomer was partially purified and reported) Mp. 114 – 117 °C; [α]^D_22_^ = −25.1 (c = 1.0, CHCl_3_); ^1^H NMR (CDCl_3_; 400 MHz) δ 4.96 (s, 1 H, OH), 4.24 – 4.20 (m, 2 H), 3.59 (td, *J* = 8, 4 Hz, 1 H), 1.84 (dt, *J* = 12, 2 Hz, 1 H), 1.70 – 0.83 (a series of m, 11 H), 1.45 (s, 3 H), 1.41 (s, 6 H, 2 CH_3_), 0.97 (s, 3 H), 0.90 (s, 3 H), 0.83 (s, 3 H) ppm; ^13^C NMR (CDCl_3_; 100 MHz) δ 107.4, 73.5, 72.8, 62.2, 55.7, 42.8, 41.5, 40.4, 37.3, 33.6, 33.3, 26.5, 26.2, 25.8, 21.7, 19.7, 18.4 (2 C), 16.1 ppm. MS (electrospray ionization), m/z 333.1 (M+Na^+^). HRMS-ESI: m/z [M + Na]^+^ calcd for C_19_H_34_NaO_3_^+^: 333.2406, found: 333.2411.

### Determination of Antifungal activity of synthetic compounds

Synthetic pure drimenol or albicanol was dissolved in DMSO (10 mg/ml as stock solution) and used for determining their antifungal activities (minimum inhibitory concentration, MIC) against various fungi according to the microdilution assay of CLSI (17). The CLSI broth dilution methods of M27-A3 for yeasts and M38-A for filamentous fungi were used to determine the susceptibility. Since our initial assay with *C. albicans* confirmed the antifungal activity of drimenol and albicanol, we extended the susceptibility assay to other pathogenic fungi including FLU resistant *C. albicans*, various species of candida, *Cryptococcus*, *Aspergillus* and a dermatophyte fungus (strains were generously provided by Dr. Ted C. White at The University of Missouri Kansas City (UMKC). *C. auris* was obtained from Dr. Baha Abdalhamid at The University of Nebraska Medical Center, Omaha NE). Briefly, yeast cells or conidia (for filamentous fungi) were suspended in RPMI 1640 medium to a final concentration of 10^5^ cfu/ml and distributed in 96-well microplate to a total volume of 100 μl/well. Drimenol or albicanol was added into the wells and a two-fold serial dilution was made. Duplicates were used for each concentration and wells with or without DMSO served as controls. Plates were incubated without shaking at 37°C for 24 - 48h for yeasts and 30°C for 4 days for filamentous fungi (*Aspergillus* sp and *Trichophyton* sp). The MIC was defined as the lowest compound concentration at which no growth occurred, as determined visually and microscopically (inverted microscope).

### Determination of *C. auris* growth inhibition by drimenol

The effect of drimenol on the growth of *C. auris* was determined by Bioscreen-C real time growth monitoring system (Oy Growth Curves Ab Ltd, Finland) as described earlier (50). Briefly, 200 µl of RPMI medium containing exponentially growing *C. auris* yeast cells (each at 0.07 OD600) were added into the honeycomb wells with or without compound (control) and measured their growth rates for 20 hours at 37°C. Compound treatment was done at two different concentrations for drimenol (50 and 60 µg/ml). The absorbance was measured at 600 nm at 30 min intervals for 24 h at 37°C with shaking for 10 s before each reading. Solvent negative control (DMSO) and FLU (60 µg/ml; antifungal drug) positive control were included in the study. The experiments were repeated at least two times with three technical replicates.

### Yeast spot assay

Yeast Peptone Dextrose (YPD) agar containing a sub-MIC concentration of drimenol (30 µg/ml) or an equal volume of DMSO was used to spot test the *C. albicans* heterozygous mutants (GRACE library (25)). Yeast suspensions of various mutants and the wild type *C. albicans* were used. Five μl of a four-fold serially diluted suspension was spotted on the agar plates and incubated at 30°C for yeast growth for 24 h, and photographed. Experiments were repeated at least three times and a representative result was shown.

### Genome-wide fitness assay

The Saccharomyces yeast deletion collection was comprised of approximately 5,900 individually bar-coded heterozygous diploid strains (HIP [haploinsufficiency profiling]) and ∼4,800 homozygous diploid strains (HOP [homozygous deletion profiling]). Pools of approximately equal strain abundance were generated by robotically pinning (S and P Robotics, Ontario, Canada) each strain (from frozen stocks) onto YPD agar plates as arrays of 384 strains/plate (21, 51, 52). After 2 days of growth at 30°C, colonies were collected from plates by flooding with YPD, and cells were adjusted to an optical density at 600 nm (OD_600_) of 2. The fitness of each strain in each experimental pool was assessed as described previously (21). The dose that resulted in 15% growth inhibition in *S*. *cerevisiae* BY4743 (the parent strain of the yeast deletion collection) was determined by analyzing dose response over the course of 16 h of growth at 30°C. Screens of the homozygous deletion collection were performed for 5 generations of growth and screens of the heterozygous deletion collection were collected after 20 generations of growth. Cells were processed as described previously (21). Genomic DNA was extracted from each sample and subjected to PCR to amplify the unique bar code identifiers. The abundance of each bar code was determined by quantifying the microarray signal as previously described (21).

*Candida albicans* pooled screens used the tn-transposon collection (23). Growth assays were performed in duplicate and samples were recovered at 20 generations of growth. Genomic DNA extraction, tag amplification, and hybridization were performed as described above.

## Supplementary data

Supplementary Table 1. *S. cerevisiae* genetic screening data

Supplemenary Table 2. *C. albicans* genetic screening data

## Acknowledgement

We thank Ted C. White, University of Missouri Kansas City (UMKC), Division of Cell Biology and Biophysics, Kansas City for various species of candida including fluconazole-resistant strains, *Cryptococcus*, *Aspergillus* and dermatophytes. Our sincere thanks to Terry Roemer, Merck Sharp and Dohme Corp, NJ for providing *C. albicans* GRACE and DBC mutant libraries. *C. auris* was obtained from Baha Abdalhamid, University of Nebraska Medical Center, Omaha NE. Kansas State University has utilized the non-clinical and pre-clinical service program offered by the National Institute of Allergy and Infectious Diseases at the Fungus Testing Laboratory, UTHSC San Antonio. Our sincere thanks to Nathan Wiederhold, Hoja Patterson, and April Torres for their assistance in testing drimenol for antifungal activity through the NIH service program. We thank Johnson Cancer Research Center, KSU for Innovative Research Award and Kansas Idea Network of Biomedical Research Excellence (K-INBRE) for CORE Facility support award to GV. CN is a CRC Tier 1 Chair in Translational Genomics and support for GG and CN is provided by the CIHR. Research reported in this publication was, in part, supported by the National Institute of General Medical Sciences of the National Institutes of Health under award number NIH R01GM128659 (to DHH). The content is solely the responsibility of the authors and does not necessarily represent the official views of the National Institutes of Health. This material was based upon work in part supported by the National Science Foundation under 1826982 (to DHH) for the purchase of a 400-MHz NMR spectrometer.

## References

1. Fuentefria AM, Pippi B, Dalla Lana DF, Donato KK, de Andrade SF. 2018. Antifungals discovery: an insight into new strategies to combat antifungal resistance. Lett Appl Microbiol 66:2–13.

2. Perfect JR. 2017. The antifungal pipeline: a reality check. Nat Rev Drug Discov 16:603–616.

3. Perlin DS, Rautemaa-Richardson R, Alastruey-Izquierdo A. 2017. The global problem of antifungal resistance: prevalence, mechanisms, and management. Lancet Infect Dis 17:e383–e392.

4. Nami S, Aghebati-Maleki A, Morovati H, Aghebati-Maleki L. 2019. Current antifungal drugs and immunotherapeutic approaches as promising strategies to treatment of fungal diseases. Biomed Pharmacother 110:857–868.

5. Nett JE, Andes DR. 2016. Antifungal Agents: Spectrum of Activity, Pharmacology, and Clinical Indications. Infect Dis Clin North Am 30:51–83.

6. Waldorf AR, Polak A. 1983. Mechanisms of action of 5-fluorocytosine. Antimicrob Agents Chemother 23:79–85.

7. Denning DW. 2002. Echinocandins: a new class of antifungal. J Antimicrob Chemother 49:889–91.

8. Roemer T, Krysan DJ. 2014. Antifungal drug development: challenges, unmet clinical needs, and new approaches. Cold Spring Harb Perspect Med 4:pii: 4/5/a019703. doi: 10.1101/cshperspect.a019703.

9. Denning DW, Hope WW. 2010. Therapy for fungal diseases: opportunities and priorities. Trends Microbiol 18:195–204.

10. Pfaller MA, Diekema DJ. 2007. Epidemiology of invasive candidiasis: a persistent public health problem. Clin Microbiol Rev 20:133–63.

11. Marie C, White TC. 2009. Genetic Basis of Antifungal Drug Resistance. Curr Fungal Infect Rep 3:163–169.

12. Sanglard D, Coste A, Ferrari S. 2009. Antifungal drug resistance mechanisms in fungal pathogens from the perspective of transcriptional gene regulation. FEMS Yeast Res 9:1029–50.

13. Hua DH, Huang X, Chen Y, Battina SK, Tamura M, Noh SK, Koo SI, Namatame I, Tomoda H, Perchellet EM, Perchellet JP. 2004. Total syntheses of (+)-chloropuupehenone and (+)-chloropuupehenol and their analogues and evaluation of their bioactivities. J Org Chem 69:6065–78.

14. Madikane VE, Bhakta S, Russell AJ, Campbell WE, Claridge TD, Elisha BG, Davies SG, Smith P, Sim E. 2007. Inhibition of mycobacterial arylamine N-acetyltransferase contributes to anti-mycobacterial activity of *Warburgia salutaris*. Bioorg Med Chem 15:3579–86.

15. Cortes M, Delgado V, Saitz C, Armstrong V. 2011. Drimenol: A versatile synthon for compounds with trans-drimane skeleton. Nat Prod Commun 6:477–90.

16. Liu Y, Liu Z, Cao X, Liu X, He H, Yang Y. 2011. Design and synthesis of pyridine-substituted itraconazole analogues with improved antifungal activities, water solubility and bioavailability. Bioorg Med Chem Lett 21:4779–83.

17. CLSI. 2008. Reference method for broth dilution antifungal susceptibility testing of yeasts and filamentous fungi; Approved Standard-Third Edition. Clinical and Laboratory Standards Institute, 940 West Valley Road, Wayne, Pa. CLSI document M27-A3 & M38-A 28.

18. Kean R, Ramage G. 2019. Combined Antifungal Resistance and Biofilm Tolerance: the Global Threat of *Candida auris*. mSphere 4:DOI: 10.1128/mSphere.00458-19.

19. de Jong AW, Hagen F. 2019. Attack, Defend and Persist: How the Fungal Pathogen *Candida auris* was Able to Emerge Globally in Healthcare Environments. Mycopathologia 184:353–365.

20. Vediyappan G, Dumontet V, Pelissier F, d’Enfert C. 2013. Gymnemic acids inhibit hyphal growth and virulence in *Candida albicans*. PLoS One 8:e74189.

21. Pierce SE, Davis RW, Nislow C, Giaever G. 2007. Genome-wide analysis of barcoded *Saccharomyces cerevisiae* gene-deletion mutants in pooled cultures. Nat Protoc 2:2958–74.

22. Xu D, Jiang B, Ketela T, Lemieux S, Veillette K, Martel N, Davison J, Sillaots S, Trosok S, Bachewich C, Bussey H, Youngman P, Roemer T. 2007. Genome-wide fitness test and mechanism-of-action studies of inhibitory compounds in *Candida albicans*. PLoS Pathog 3:e92.

23. Oh J, Fung E, Schlecht U, Davis RW, Giaever G, St Onge RP, Deutschbauer A, Nislow C. 2010. Gene annotation and drug target discovery in *Candida albicans* with a tagged transposon mutant collection. PLoS Pathog 6:e1001140.

24. Chen J, Zhou S, Wang Q, Chen X, Pan T, Liu H. 2000. Crk1, a novel Cdc2-related protein kinase, is required for hyphal development and virulence in *Candida albicans*. Mol Cell Biol 20:8696–708.

25. Roemer T, Jiang B, Davison J, Ketela T, Veillette K, Breton A, Tandia F, Linteau A, Sillaots S, Marta C, Martel N, Veronneau S, Lemieux S, Kauffman S, Becker J, Storms R, Boone C, Bussey H. 2003. Large-scale essential gene identification in *Candida albicans* and applications to antifungal drug discovery. Mol Microbiol 50:167–81.

26. An T, Liu Y, Gourguechon S, Wang CC, Li Z. 2018. CDK Phosphorylation of Translation Initiation Factors Couples Protein Translation with Cell-Cycle Transition. Cell Rep 25:3204–3214 e5.

27. CGD. 2019. Candida Genome Database, http://www.candidagenome.org/cgi-bin/locus.pl?locus=Sec21&organism=C_albicans_SC5314.

28. CGD. 2019. Candida Genome Database, (http://www.candidagenome.org/cgi-bin/locus.pl?locus=ret2&organism=C_albicans_SC5314.

29. Higuchi Y. 2015. Initial fungal effector production is mediated by early endosome motility. Commun Integr Biol 8:e1025187.

30. Bielska E, Higuchi Y, Schuster M, Steinberg N, Kilaru S, Talbot NJ, Steinberg G. 2014. Long-distance endosome trafficking drives fungal effector production during plant infection. Nat Commun 5:5097.

31. Wernimont AK, Dong A, Lew J, Lin YH, Hassanali A, Arrowsmith CH, Edwards AM, Weigelt J, Sundstrom M, Bochkarev A, Hui R, Artz JD, (Structural Genomics Consortium). 2007. Cryptosporidium parvum cyclin-dependent kinase cgd5_2510 with indirubin 3’-monoxime bound. https://wwwrcsborg/structure/2qkr (database release).

32. Trott O, Olson AJ. 2010. AutoDock Vina: improving the speed and accuracy of docking with a new scoring function, efficient optimization, and multithreading. J Comput Chem 31:455–61.

33. Pfaller MA, Diekema DJ. 2010. Epidemiology of invasive mycoses in North America. Crit Rev Microbiol 36:1–53.

34. Pappas PG. 2006. Invasive candidiasis. Infect Dis Clin North Am 20:485–506.

35. Pfaller MA, Diekema DJ, Rinaldi MG, Barnes R, Hu B, Veselov AV, Tiraboschi N, Nagy E, Gibbs DL. 2005. Results from the ARTEMIS DISK Global Antifungal Surveillance Study: a 6.5-year analysis of susceptibilities of Candida and other yeast species to fluconazole and voriconazole by standardized disk diffusion testing. J Clin Microbiol 43:5848–59.

36. Park BJ, Wannemuehler KA, Marston BJ, Govender N, Pappas PG, Chiller TM. 2009. Estimation of the current global burden of cryptococcal meningitis among persons living with HIV/AIDS. AIDS 23:525–30.

37. Byrnes EJ, 3rd, Li W, Lewit Y, Ma H, Voelz K, Ren P, Carter DA, Chaturvedi V, Bildfell RJ, May RC, Heitman J. 2010. Emergence and pathogenicity of highly virulent *Cryptococcus gattii* genotypes in the northwest United States. PLoS Pathog 6:e1000850.

38. Kidd SE, Hagen F, Tscharke RL, Huynh M, Bartlett KH, Fyfe M, Macdougall L, Boekhout T, Kwon-Chung KJ, Meyer W. 2004. A rare genotype of *Cryptococcus gattii* caused the cryptococcosis outbreak on Vancouver Island (British Columbia, Canada). Proc Natl Acad Sci U S A 101:17258–63.

39. NPR-News. 2010. Fungal disease spreads through Pacific Northwest; http://www.npr.org/templates/story/story.php?storyId=126198896.

40. Alangaden GJ. 2011. Nosocomial fungal infections: epidemiology, infection control, and prevention. Infect Dis Clin North Am 25:201–25.

41. Neofytos D, Horn D, Anaissie E, Steinbach W, Olyaei A, Fishman J, Pfaller M, Chang C, Webster K, Marr K. 2009. Epidemiology and outcome of invasive fungal infection in adult hematopoietic stem cell transplant recipients: analysis of Multicenter Prospective Antifungal Therapy (PATH) Alliance registry. Clin Infect Dis 48:265–73.

42. Achterman RR, Smith AR, Oliver BG, White TC. 2011. Sequenced dermatophyte strains: growth rate, conidiation, drug susceptibilities, and virulence in an invertebrate model. Fungal Genet Biol 48:335–41.

43. Butler MS. 2008. Natural products to drugs: natural product-derived compounds in clinical trials. Nat Prod Rep 25:475–516.

44. Jansen BJ, de Groot A. 2004. Occurrence, biological activity and synthesis of drimane sesquiterpenoids. Nat Prod Rep 21:449–77.

45. Toyota M, Ooiso Y, Kusuyama T, Asakawa Y. 1994. Drimane-type sesquiterpenoids from the liverwort *Diplophyllum serrulatum*. Phytochemistry 35:1263–1265.

46. Robles-Kelly C, Rubio J, Thomas M, Sedan C, Martinez R, Olea AF, Carrasco H, Taborga L, Silva-Moreno E. 2017. Effect of drimenol and synthetic derivatives on growth and germination of *Botrytis cinerea*: Evaluation of possible mechanism of action. Pestic Biochem Physiol 141:50–56.

47. Giaever G, Shoemaker DD, Jones TW, Liang H, Winzeler EA, Astromoff A, Davis RW. 1999. Genomic profiling of drug sensitivities via induced haploinsufficiency. Nat Genet 21:278–83.

48. Kubo I, Fujita K, Lee SH. 2001. Antifungal mechanism of polygodial. J Agric Food Chem 49:1607–11.

49. Fujita K, Kubo I. 2005. Multifunctional action of antifungal polygodial against *Saccharomyces cerevisiae*: involvement of pyrrole formation on cell surface in antifungal action. Bioorg Med Chem 13:6742–7.

50. Veerapandian R, Vediyappan G. 2019. Gymnemic acids inhibit adhesive nanofibrillar mediated *Streptococcus gordonii-Candida albicans* mono-species and dual-species biofilms. Front Microbiol 10: 2328.

51. Mor V, Rella A, Farnoud AM, Singh A, Munshi M, Bryan A, Naseem S, Konopka JB, Ojima I, Bullesbach E, Ashbaugh A, Linke MJ, Cushion M, Collins M, Ananthula HK, Sallans L, Desai PB, Wiederhold NP, Fothergill AW, Kirkpatrick WR, Patterson T, Wong LH, Sinha S, Giaever G, Nislow C, Flaherty P, Pan X, Cesar GV, de Melo Tavares P, Frases S, Miranda K, Rodrigues ML, Luberto C, Nimrichter L, Del Poeta M. 2015. Identification of a New Class of Antifungals Targeting the Synthesis of Fungal Sphingolipids. MBio 6:e00647.

52. Proctor M, Urbanus ML, Fung EL, Jaramillo DF, Davis RW, Nislow C, Giaever G. 2011. The automated cell: compound and environment screening system (ACCESS) for chemogenomic screening. Methods Mol Biol 759:239–69.

